# SANS serif: alignment-free, whole-genome based phylogenetic reconstruction

**DOI:** 10.1101/2020.12.31.424643

**Authors:** Andreas Rempel, Roland Wittler

## Abstract

**Summary:** SANS serif is a novel software for alignment-free, whole-genome based phylogeny estimation that follows a pangenomic approach to efficiently calculate a set of splits in a phylogenetic tree or network.

**Availability and Implementation:** Implemented in C++ and supported on Linux, MacOS, and Windows. The source code is freely available for download at https://gitlab.ub.uni-bielefeld.de/gi/sans.

## 1 Introduction

In computational pangenomics and phylogenomics, a major challenge is the memory- and time-efficient analysis of multiple genomes in parallel. Conventional approaches for the reconstruction of phylogenetic trees are based on the alignment of specific sequences such as marker genes. However, the problem of multiple sequence alignment is complex and practically infeasible for large-scale data. Whole-genome approaches do neither require the identification of marker genes nor expensive alignments, but they usually perform a quadratic number of sequence comparisons (in terms of *k*-mers or other patterns) to obtain pairwise distances. This leads to a runtime that increases quadratically with the number of input sequences and is not suitable for projects comprising a large number of genomes.

We present the software SANS serif, which is based on a whole-genome approach and is both alignment- and reference-free. The command line tool accepts both assembled genomes and raw reads as input and calculates a set of splits that can be visualized as a phylogenetic tree or network using existing tools such as SplitsTree (Huson and Bryant, 2006). Instead of computing pairwise distances, the tool follows a pangenomic approach that does not require a quadratic number of sequence comparisons. The evaluation of our previous implementation SANS (Wittler, 2020) already showed promising results and revealed that our method is significantly faster and more memory-efficient than other whole-genome based approaches. The new version SANS serif is a standalone re-implementation that does not rely on 3rd-party libraries, introduces several new features, and improves the performance of our method even further, reducing the runtime and memory usage to only about 20 % compared to the original implementation.

## 2 Features and approach

The general idea of our method is to determine the evolutionary relationship of a set of genomes based on the similarity of their whole sequences. Common sequence segments that are shared by a subset of genomes are used as an indicator that these genomes lie closer together in the phylogeny and should be separated from the set of all other genomes that do not share these segments. Each pair of such two sets forms a phylogenetic split, based on the concept of Bandelt and Dress (1992), and the lengths of the concerned segments contribute to the weight of the split, i.e., the length of the edge separating these two sets in the phylogeny.

SANS serif accepts a list of multiple FASTA or FASTQ files containing complete genomes, assembled contigs, or raw reads as input. In addition, the program offers the option to import a colored de Bruijn graph generated with the software Bifrost (Holley and Melsted, 2019). The lengths of the common sequences are counted in terms of *k*-mers, i.e., overlapping substrings of length *k*. The tool is capable of handling ambiguous IUPAC characters such as N’s, replacing these with the corresponding DNA bases, considering all possibilities. The output of the program is a tab-separated text file listing all calculated splits ordered by weight (or NEWICK format if applicable). Representing a phylogeny by a list of splits has the advantage that it allows to capture ambiguous signals in the input data that may arise, e.g., due to horizontal gene transfers and would be lost in a conventional phylogenetic tree. In these cases, the output corresponds to a phylogenetic network with the ambiguous signals appearing as parallel edges, as can be seen in Figure 1C. Several filter options allow to limit the output to a fixed number of most significant splits, to reduce the complexity of the network, or to calculate a subset of the splits representing a tree.

**Figure 1:**
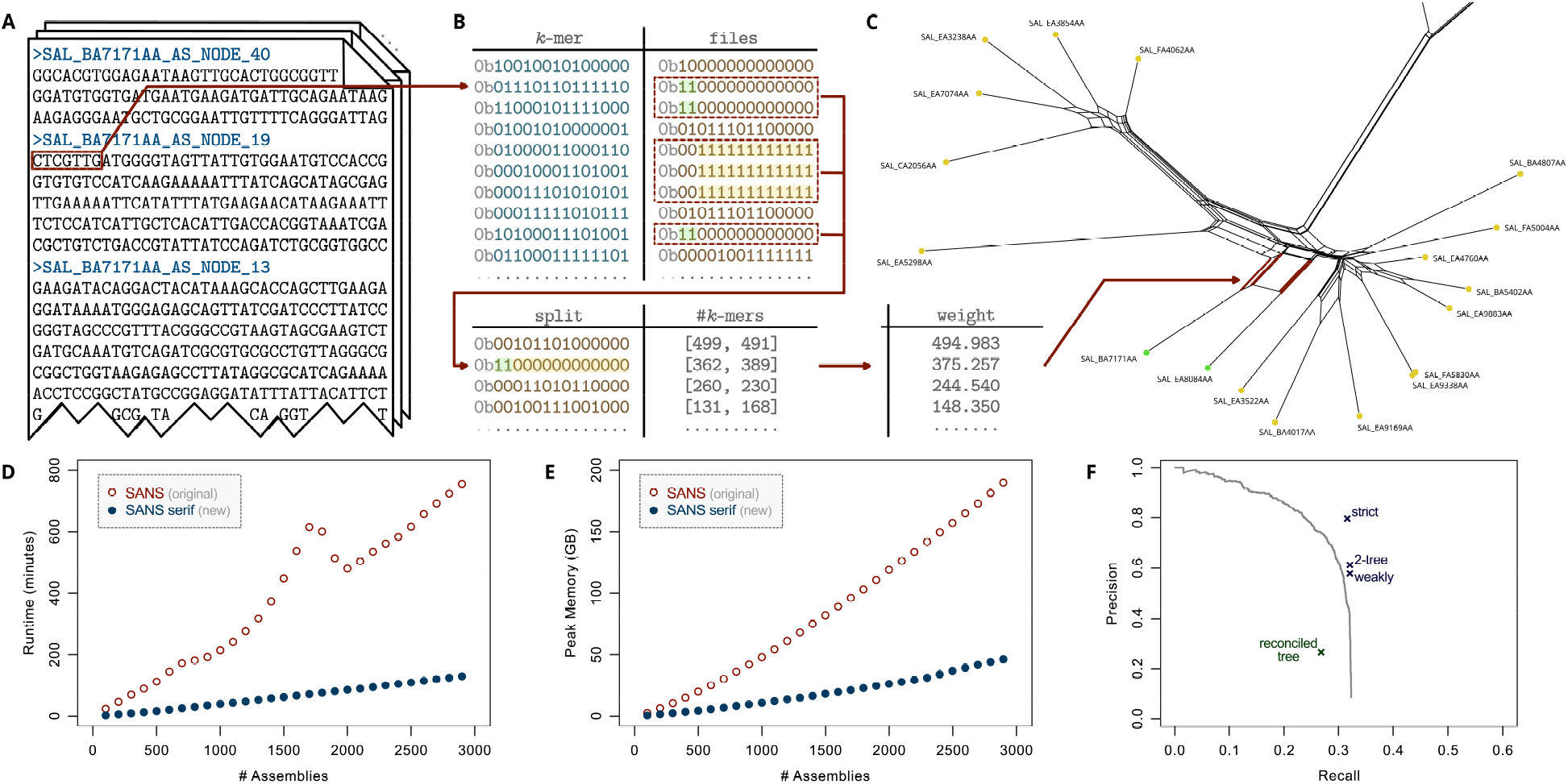
SANS serif methodology and evaluation. (A) One sequence file (FASTA/FASTQ) per genome/assembly is provided as input and *k*-mers are extracted and stored in a hash table. (B) For each *k*-mer, a bit vector is stored to encode its presence or absence: a one (or zero) at position *i* indicates its presence (resp. absence) in the *i*-th input file. Presence/absence patterns are combined to *splits* and stored in a second hash table together with two counts: the number of *k*-mers having that pattern and the number of *k*-mers having the complementary pattern. (C) Both counts are combined to an overall weight per split. Splits can be filtered and visualized as a network. As an example, a subnet of a weakly compatible split set of the *Salmonella* data set (Zhou *et al.*, 2018) is shown. Splits are represented by parallel edges. (D, E) Runtime and peak memory usage of SANS on the *Salmonella* data set on a single 2 GHz processor. For random subsamples of *n* assemblies and *k*=21, the 10*n* highest weighting splits were output. Values were averaged over processing three random subsamples each. (F) Accuracy w.r.t. the reference tree. Precision: (number of called splits also in the reference tree) / (number of all called splits), Recall: (number of reference splits also in the call set) / (number of all reference splits). Trivial splits, i.e., splits separating single leaves, are not considered. Filters: only considering the *x* highest weighting splits for increasing *x* (line), greedily extracting a subset that corresponds to a tree (strict), additionally keeping a second subset that corresponds to a tree (2-tree), greedily extracting a subset that is weakly compatible (weakly). For comparison, the accuracy of a reconciled tree of 3002 core genes (Zhou *et al.*, 2018, Fig. 2A, supertree 2) is included, emphasizing the complexity of the data set and the sensibility of the applied measure.

## 3 Implementation

SANS serif is implemented in C++ and runs on all major platforms. In contrast to the previous version, the program can be compiled and executed without any dependencies. The tool owes its good performance mainly to the way that the sequences are processed and stored in memory. Each input file is processed by moving a window of a user-defined length *k* over the sequences and extracting the *k*-mers, as illustrated in Figure 1A. The program uses a hash table to store each *k*-mer as a key and a marker for the set of input files in which the *k*-mer is present as a value, updating the marker whenever the *k*-mer is observed in a new sequence. Both the *k*-mers and the markers are encoded as bit vectors in order to reduce the amount of memory and enable fast bitwise comparisons and set operations. After all files have been processed, the table is scanned once to count the *k*-mers that belong to the same subset of input files. Each pair of complementary subsets is combined to one split, accumulating the corresponding two numbers of *k*-mers separately, as illustrated in Figure 1B. To this end, a second hash table is used with the split, represented by the bit vector of the smaller of the two subsets, as a key and the two *k*-mer numbers as a value. The overall weight of each split is calculated using the geometric mean of both numbers and the selected filter is applied.

## 4 Results

We evaluated the performance of our program on a data set comprising 2964 genomes of *Salmonella enterica* subspecies *enterica* (Zhou *et al.*, 2018). Figures 1D and E show the runtime and memory usage of SANS serif compared to the original version of SANS depending on the number of input genomes. Supplementary Figures 1 and 2 additionally show the runtime and memory usage in dependence on the chosen *k*-mer length. An earlier evaluation (Wittler, 2020) already revealed that SANS demands significantly less resources than comparable tools. The new version SANS serif manages to reduce the runtime and memory usage even further by about 80 %, processing the complete data set on a single 2 GHz processor in almost 2 hours using less than 50 GB of RAM (cf. Supplementary Table 1). Figure 1F shows the accuracy of the calculated split sets and filters. To exemplify the ability to handle ambiguous IUPAC characters, we simulated a data set of 100 genomes and incorporated N’s at random positions. The accuracy dropped significantly when the affected *k*-mers were skipped, whereas the new functionality to process these characters allowed to almost recover the original accuracy (cf. Supplementary Table 2).

## 5 Conclusion

SANS serif is a novel software that allows whole-genome based phylogeny estimation with an unprecedented performance. We hope that our tool will open the way to phylogenetic analysis for data sets where it was previously not possible. A documentation listing all required and optional parameters as well as some quick start examples can be found on the project website.

## Acknowledgements

The authors thank Jens Stoye for helpful advice and acknowledge the Bielefeld-Gießen Center for Microbial Bioinformatics (BiGi) of the BMBF-funded German network for bioinformatics infrastructure (de.NBI) [grant 031A533] for provision of compute resources and general support.

## Funding

This project has received funding from the European Union’s Horizon 2020 research and innovation programme under the Marie Skłodowska-Curie agreement [grant number 872539].

## Notes

https://gitlab.ub.uni-bielefeld.de/gi/sans

### Competing Interest Statement

The authors have declared no competing interest.

