## Supplementary Material for "SANS serif: alignment-free, whole-genome based phylogenetic reconstruction"

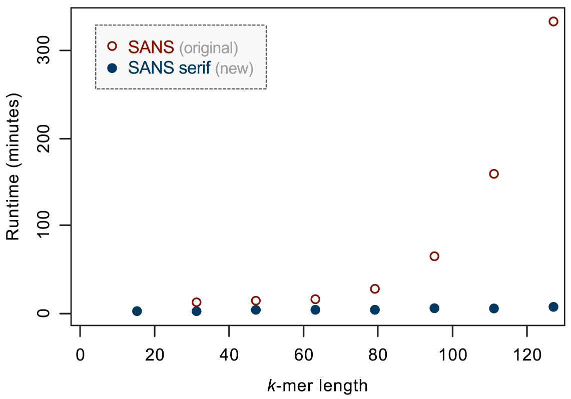


Figure 1: Runtime of SANS evaluated for different $k$-mer lengths. For random subsamples of 100 assemblies from the *Salmonella* data set, the 1000 highest weighting splits were output. Values were averaged over processing three random subsamples each.

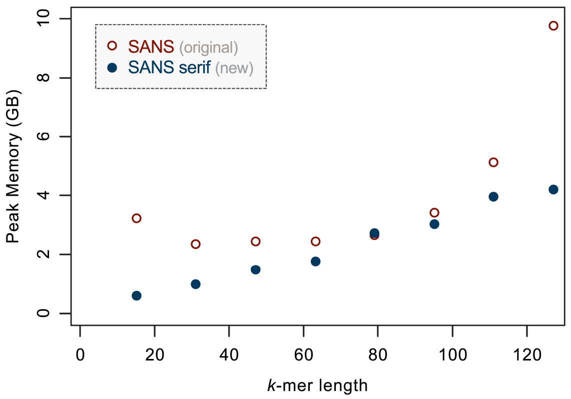


Figure 2: Peak memory usage of SANS evaluated for different $k$-mer lengths. For random subsamples of 100 assemblies from the *Salmonella* data set, the 1000 highest weighting splits were output. Values were averaged over processing three random subsamples.

Table 1: Runtime and peak memory usage of SANS serif on the complete *Salmonella* data set evaluated for different parameters: processing the sequence files directly (-input), processing a colored de Bruijn graph generated from the sequences using Bifrost (-graph), processing a list of splits and applying different filters (-filter), processing the sequence files considering IUPAC characters (-iupac). Values were averaged over three different runs.

| Parameter | | Runtime (min) | Memory (GB) |
|---|---|---|---|
| -input | | 132.71 | 47.57 |
| -graph | | 69.08 | 51.28 |
| -filter | *strict* | 0.06 | 0.01 |
| | *2-tree* | 0.06 | 0.01 |
| | *weakly* | 467.89 | 0.01 |
| -iupac | | 153.29 | 47.59 |

Table 2: Effect of N's in the input sequences. A phylogeny with 100 leaf genomes of length ~96 kb and an evolutionary distance of 5 PAM to the root has been simulated with ALF (Dalquen *et al.*, 2012). (See software repository for the complete parameter setting.) DNA characters in the genomes were randomly substituted by N's at a rate of 0.1 % and the sequences were processed by SANS serif before and after the substitution with $k$-mers containing N's being skipped (default) or N's replaced by all possible bases (parameter -iupac, $x$=16). For the obtained tree-filtered split sets, precision and recall (cf. Figure 1) as well as weighted precision (total weight of called splits also in the reference tree) / (total weight of all called splits) and weighted recall (total weight of reference splits also in the call set) / (total weight of all reference splits) have been computed.

| | original | with 0.1 % N's | |
|---|---|---|---|
| | | skipped | replaced |
| *unweighted* | | | |
| precision | 0.90 | 0.63 | 0.78 |
| recall | 0.70 | 0.38 | 0.58 |
| *weighted* | | | |
| precision | 0.98 | 0.95 | 0.98 |
| recall | 0.88 | 0.68 | 0.88 |
